# Preservation of three-dimensional spatial structure in the gut microbiome

**DOI:** 10.1101/175224

**Authors:** Yuko Hasegawa, Jessica L. Mark Welch, Blair J. Rossetti, Gary G. Borisy

**Affiliations:** Marine Biological Laboratory, Woods Hole, Massachusetts, United States of America; The Forsyth Institute, Cambridge, Massachusetts, United States of America

## Abstract

Preservation of three-dimensional structure in the gut is necessary in order to analyze the spatial organization of the gut microbiota and gut luminal contents. In this study, we evaluated preparation methods for mouse gut with the goal of preserving micron-scale spatial structure while performing fluorescence imaging assays. Our evaluation of embedding methods showed that commonly used media such as Tissue-Tek Optimal Cutting Temperature (OCT) compound, paraffin, and polyester waxes resulted in redistribution of luminal contents. By contrast, a hydrophilic methacrylate resin, Technovit H8100, preserved three-dimensional organization. Our mouse intestinal preparation protocol optimized using the Technovit H8100 embedding method was compatible with microbial fluorescence *in situ* hybridization (FISH) and other labeling techniques, including immunostaining and staining with both wheat germ agglutinin (WGA) and 4’,6-diamidino-2-phenylindole (DAPI). Mucus labeling patterns of the samples fixed with paraformaldehyde (PFA) and Carnoy’s fixative were comparable. The protocol optimized in this study enabled simultaneous visualization of micron-scale spatial patterns formed by microbial cells in the mouse intestines along with biogeographical landmarks such as host-derived mucus and food particles.

## Introduction

Preservation of spatial structure in intestinal samples is crucial for investigating spatial organization of the gut microbiota relative to mucins, host tissue, and food particles. Yet selecting a protocol for intestinal sample preparation is made difficult by conflicting recommendations in the literature. Several authors report that fixation in aqueous fixatives such as formaldehyde allows dissolution or dispersal of the mucus layer and suggest that fixation in the non-aqueous Carnoy or methacarn solution is essential for mucus preservation (Swidsinski 2005, Johansson & Hansson 2012, Earle et al. 2015). Thus, a widely used protocol is to process gut samples by Carnoy fixation followed by paraffin embedding and sectioning. Other authors, however, report that the use of organic solvents during the paraffin clearing step results in extraction of portions of the mucus layer and instead recommend freezing in Optimal Cutting Temperature (OCT) compound and cryosectioning followed by fixation in formalin for optimal preservation of mucus (Cohen et al. 2012). An alternative method, embedment in acrylic resin, has long been used for electron microscopy and is increasingly being applied to samples to be examined using light microscopy and fluorescence *in situ* hybridization (FISH) (Moter et al. 1998, Moter & Göbel 2000, De Jonge et al. 2005, Heimesaat et al. 2006, Zijnge et al. 2010, Schimak et al. 2016). However, in no case has a thorough investigation been carried out to determine if the protocols adequately preserve the three-dimensional organization of the gut luminal contents.

In this study, we compared protocols for preparation of intestinal sections with the goal of preserving the micron-scale spatial arrangement of microbial cells and other luminal contents, including food particles and host-derived mucus. We compared preservation of three-dimensional structure using confocal microscopy of four different embedding media: OCT compound; paraffin wax; polyester wax; and glycol methacrylate resin. We then evaluated applicability of microbial FISH and two mucus labeling methods to simultaneously visualize microbial cells and host-derived mucus in intestinal sections following both Carnoy and paraformaldehyde (PFA) fixation.

## Materials and methods

### Mouse intestinal samples

Segments of colon were collected from nu/nu mice (courtesy of Dr. Shanta M. Messerli, Marine Biological Laboratory) euthanized for unrelated experiments. Protocols for animal care, handling, and “tissue sharing” were approved by the Institutional Animal Care and Use Committee of the Marine Biological Laboratory. Additional samples from gnotobiotic C57BL/6J mice were provided by Dr. Nathan McNulty and Dr. Jeffrey I. Gordon, Washington University in St. Louis. Intestinal samples were cut into 5- to 10-mm long pieces using a razor blade and subsequently processed by one of the following methods.

### Methacrylate embedding with Carnoy fixation

Samples were placed directly into ice-cold Carnoy solution (60% ethanol, 30% chloroform, 10% glacial acetic acid) for 2 hours, then rinsed with several changes of 100% ethanol and stored in 100% ethanol at −20°C until embedding. For embedding in Technovit H8100 methacrylate resin, the ethanol was removed and samples were infiltrated with several changes of Technovit H8100 infiltration solution prepared as directed by the manufacturer under vacuum at 4°C with gentle agitation, with the last infiltration step proceeding overnight. Samples were then transferred into BEEM capsules filled with embedding solution prepared as directed by the manufacturer and allowed to harden at 4°C overnight.

### Methacrylate embedding with PFA fixation

Samples were gently coated with molten 0.5% low melting point agarose, placed at 4°C to allow the agarose to harden, then fixed in 2% PFA in phosphate-buffered saline (PBS) for 12 hours at 4°C. Samples were briefly washed with 1X PBS and deionized water, then were again coated in molten 0.5% agarose and placed at 4°C to allow the agarose to harden. Excess agarose was trimmed before Technovit H8100 embedding. Samples were dehydrated in acetone for 1 hour at 4 °C and were then placed in infiltration solution prepared as directed by the manufacturer with several changes of infiltration solution for a total of 12 hours. Samples were then transferred into Eppendorf tubes filled with embedding solution prepared as directed by the manufacturer and allowed to harden for 12 hours at 4 °C.

### Sectioning of methacrylate blocks

Methacrylate-embedded intestines were sectioned to 5 μm thickness using a microtome (Sorvall JB-4, Dupont Instrument). Sections were cut with glass knives or tungsten carbide triangle knives and were sectioned dry and transferred onto a drop of water on a slide. Sections were dried on a warm plate and then subjected to fluorescent labeling experiments.

### Paraffin embedding with Carnoy fixation

Samples were immersed in ice-cold Carnoy solution for 2 hours, then rinsed in several changes of 100% ethanol and stored in 100% ethanol at −20°C until embedding. For embedding, samples were immersed in 3 changes of xylene for one hour each, then immersed in 3 changes of molten paraffin wax (Paraplast, Electron Microscopy Sciences) at 56-58°C for one hour each. Blocks were allowed to harden at room temperature. Sections were cut to 15 μm thickness and were floated on a water bath at 40-45°C, then transferred to slides, dried, and stored at room temperature. In preparation for staining, slides were deparaffinized by heating at 60°C for 10 minutes followed by immersion in 4 changes of xylene for 2.5 minutes each, then in 2 changes of 100% ethanol for 3 minutes each, then rehydrated through 95% and 75% ethanol for 1 minute each and immersed in 900 mM NaCl, 20 mM Tris pH 7.5 in preparation for FISH.

### Polyester embedding with Carnoy fixation

Samples were immersed in ice-cold Carnoy solution for 2 hours, then rinsed in several changes of 100% ethanol and stored in 100% ethanol at −20°C until embedding. For embedding, samples were immersed in a 1:1 mixture of molten polyester wax (polyethylene glycol distearate PEG-400 DS, HallStar Company, Stow, OH) and ethanol at 45°C for 30 minutes, then in 3 changes of 100% molten polyester wax at 45°C for 1 hour each. Blocks were allowed to harden at room temperature. Sections were cut to 15 μm thickness and were transferred to a subbed slide, stretched using a drop of 2% PFA, dried, and stored at room temperature. In preparation for staining, slides were incubated in 2 changes of 100% ethanol for 5 minutes each, then in 95% and 75% ethanol for 5 minutes each, then immersed in 900 mM NaCl, 20 mM Tris pH 7.5 in preparation for FISH.

### Cryosectioning

Intestinal samples were immersed in OCT compound (Tissue-Tek, Sakura Finetek USA, Inc. Torrance, CA), snap-frozen in liquid nitrogen and stored at −80 °C until sectioning. Frozen samples were sectioned to 12.5 μm thickness at −25 °C using a cryostat (Microm HM 505N) and then placed on Superfrost Plus microscope slides (Cat # 4951PLUS-001, Erie Scientific Company, Portsmouth, NH). To prevent the loss of intestinal contents as well as sections themselves during fixation and labeling, sections were coated with agarose by touching the surface of the slide to molten 0.5% low melting point agarose (IB70051, IBI Scientific, Peosta, IA). Sections were then fixed in 2% PFA in 1x PBS for 12 hours at 4°C, then briefly washed with 1 X PBS and distilled water to remove PFA. Slides were then dipped for 3 minutes each into 50%, 80%, and 96% (v/v) ethanol to dehydrate the intestinal sections.

### FISH

FISH targeting bacterial cells was performed using the oligonucleotide probe Eub338 (Amann et al. 1990) custom synthesized and 5’ labeled with Rhodamine Red X (Thermo-Fisher Inc., Waltham, MA). Sections were incubated in a hybridization buffer of 0.9 M NaCl, 0.02 M Tris pH 7.5, 0.01% SDS, 20% HiDi formamide (Applied Biosystems) and 2 μM probe at 46°C for six hours. After hybridization, samples were washed at 48°C for 15 minutes in a large excess of wash buffer (0.215 M NaCl, 0.02 M Tris pH 7.5, 0.005 M EDTA). Additional FISH on sections of gnotobiotic colon was carried out in the same way except using three oligonucleotide probes: Eub338 5’ labeled with Alexa Fluor 647, Bthe577 specific for *Bacteroides thetaiotaomicron* (Mark Welch et al., submitted) 5’ labeled with Rhodamine Red X and Erec1259 specific for *Eubacterium rectale* (Mark Welch et al., submitted) 5’ labeled with Alexa Fluor 594.

### Immunostaining

Sections on microscope slides were treated with a blocking solution (2% goat serum; 1% bovine serum albumin; 0.2% Triton X-100; 0.05% Tween 20) for 1 hour at room temperature. Sections were then incubated with 1:50 dilution of an anti-mouse colonic mucin primary antibody (anti-MCM, a gift of Dr. Ingrid B. Renes, Erasmus MC-Josephine Nefkens Institute) diluted in the blocking solution for 12 hours at 4 °C. After incubation, intestinal sections were rinsed for 3 minutes each in fresh 1X PBS, treated with blocking solution for 1 hour at room temperature, and then incubated with a 1:1000 dilution of the Alexa Fluor 633 goat anti-rabbit IgG (Cat# A21070, Invitrogen, Carlsbad, CA) in blocking solution and rinsed for 3 minutes each in fresh 1X PBS.

### Fluorescent staining with DAPI and WGA

Sections were stained with 1 μg/ml 4’,6-diamidino-2-phenylindole (DAPI) and 20 or 40 μg/ml wheat germ agglutinin (WGA) Alexa Fluor 680 conjugate (Invitrogen Inc., Carlsbad, CA) in 1X PBS for 15 minutes at room temperature, then incubated 2 x 3 minutes in wash buffer (215 mM NaCl, 20 mM Tris pH 7.5, 5 mM EDTA). Slides were then dipped in water, drained, and air-dried, or were dipped for 3 minutes each into 50%, 80%, and 96% (v/v) ethanol and then air-dried. Slides were mounted with ProLong Gold antifade reagent (Cat # P36934, Invitrogen Inc., Carlsbad, CA) and placed in a dark environment at room temperature for at least 24 hours until the mounting medium solidified. Images were acquired with a Zeiss LSM 510, 710 or 780 confocal microscope or a widefield microscope, the Axio Imager.Z2 (Carl Zeiss, Thornwood, NY). When microbial cells and mucus were visualized on the same Technovit H8100-embedded intestinal sections, FISH was performed prior to mucus staining.

## Results

### Retention of luminal contents

Following standard procedures, we first attempted to immobilize, retain and visualize luminal contents by a cryo-freezing and cryosectioning procedure. We immersed pieces of freshly dissected intestine in OCT compound, snap-froze them in liquid nitrogen and stored them at −80°C until cryosectioning at −25°C. However, in initial experiments, we found that a large fraction of the luminal contents, including both food particles and bacteria, were apparently missing or lost from the sample sections (Fig 1 top row). Partial or complete loss occurred irrespective of whether the OCT-embedded cryosections were fixed with PFA or Carnoy’s fixative or with both. Since most of the luminal contents are normally in physiological transit through the gut, they are intrinsically not adherent to the gut wall. We inferred that the luminal contents could have been physically lost from the sections even though the material had been subject to fixation. Loss could have occurred prior to chemical fixation, during the fixation process itself, washing steps or during the subsequent staining.

**Figure 1.**
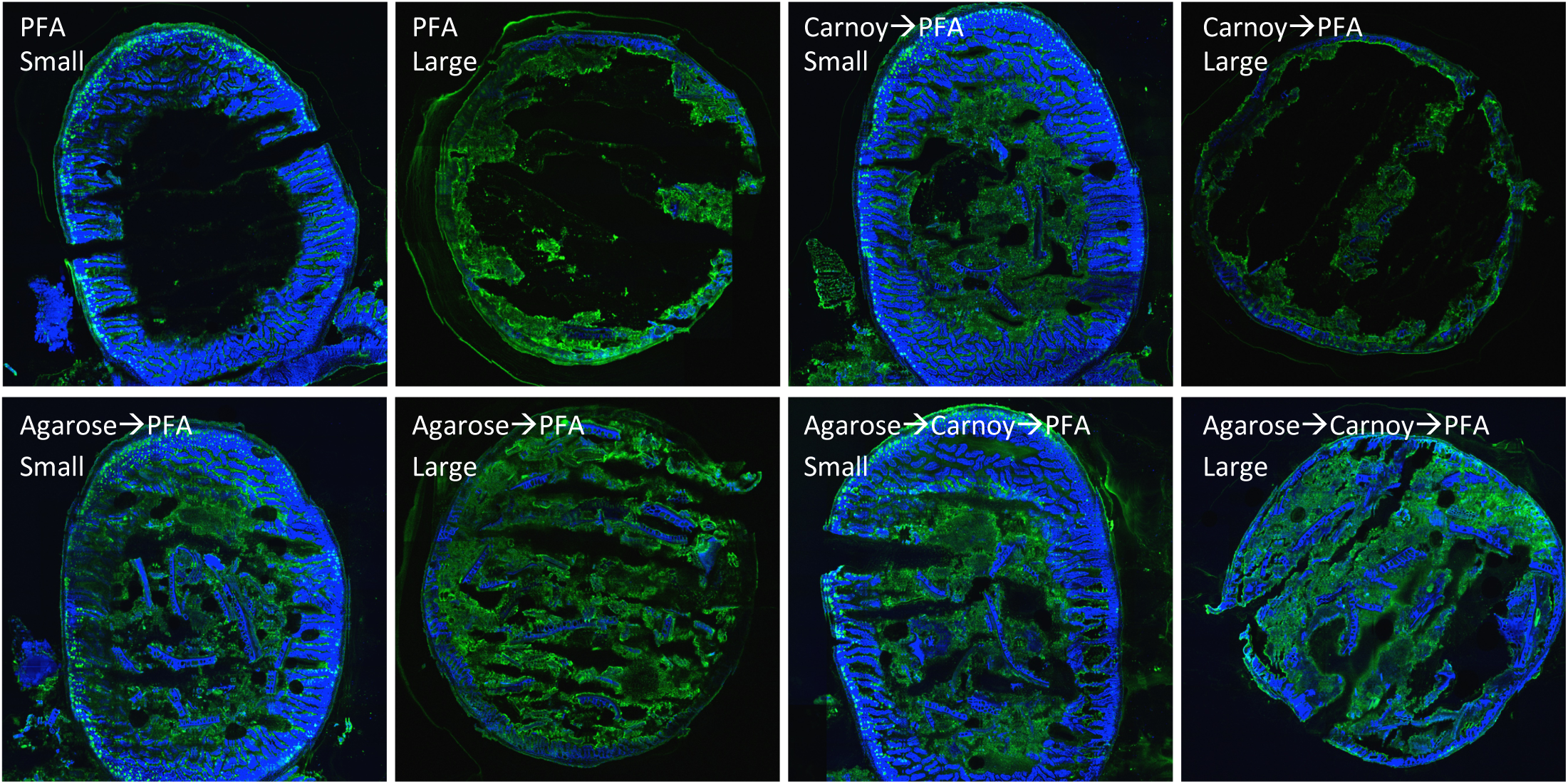
Agarose coating reduces loss of material from cryosections. Cryosections of small or large intestine were fixed with PFA with or without prior immersion in Carnoy solution (top row). For comparison, cryosections were coated with 1% low-melting point agarose and then subjected to the same fixation procedures. All samples were then stained with DAPI (blue) and Alexa fluor 488-labeled wheat germ agglutinin (green). For this test, dual-taxa colonized large and small intestinal samples were used.

Since sample cryosections are normally warmed to 4°C for post-sectioning fixation, cryo-immobilization would have been lost during the warming. Consequently, we explored the use of a hydrogel to provide immobilization before subjecting the cryosections to any further procedures. Nascent sections were exposed to a thin layer of low melting point agarose immediately after cryosectioning. After gelling, the sections were then fixed and processed. This simple agarose embedding step retained luminal contents and showed little or no sign of loss of material (Fig 1 bottom row).

For preparation of colon samples for imaging, we explored four sectioning procedures: cryosectioning with post-sectioning fixation, and structure preservation by chemical fixation prior to embedment and sectioning, for which we explored three different embedment materials—paraffin wax, polyester wax, and glycol methacrylate plastic (Technovit H8100). Either freshly dissected intestinal segments or segments previously snap-frozen were fixed at 4°C prior to embedment and sectioning at room temperature. In this procedure, two agarose hydrogel steps were incorporated into the protocol. First, molten agarose was applied to the ends of the intestinal segments and allowed to harden to prevent loss of material from the ends. Then, the tissue segments were fixed (in either PFA or Carnoy’s fixative) after which they were immersed in molten agarose and allowed to harden in preparation for cutting into smaller pieces for embedment. By these methods (Fig 2), luminal contents, including both food and bacteria, were retained and could be readily visualized in conventional sections stained with DAPI or prepared for fluorescence *in situ* hybridization.

**Figure 2.**
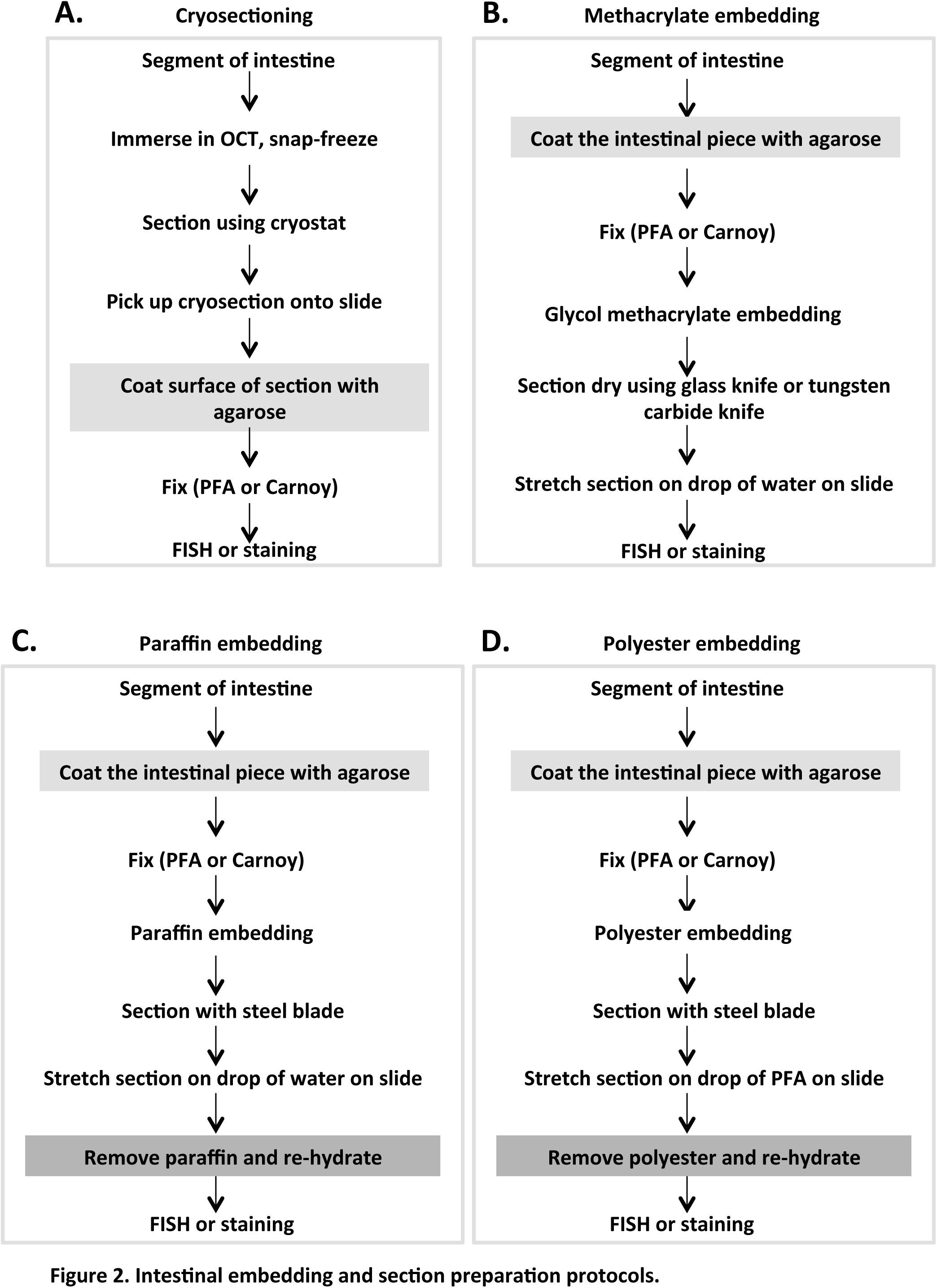
Intestinal embedding and section preparation protocols.

### Evaluation of 3D preservation

Having established several procedures for retaining and visualizing intestinal luminal contents, the next issue to be investigated was the degree to which the three-dimensional distribution of material was preserved. To evaluate this question, confocal *z*-stack images were acquired for each of the preparative procedures. Although conventional two-dimensional images acquired in the *xy*-plane showed apparently similar features for each of the procedures (Fig 3), analysis in the *z*-direction revealed important differences. As demonstrated by profile images in the *xz*- or *yz*-planes (Fig 4), only for embedment in Technovit H8100 were the luminal contents evenly distributed throughout the depth of the section. For cryofrozen, cryosectioned, and post-fixed samples and for prefixed paraffin or polyester embedded samples, luminal contents were not evenly distributed through the section. Analysis of the *xz*- and *yz*-planes showed that much of the luminal material was at the bottom of the section, the surface adjacent to the microscope slide. This uneven and bottom-heavy distribution suggests that luminal contents became redistributed by collapsing onto the slide during preparation, or that the adhesive surface of the slide retained only the material immediately adjacent to it, while material not adjacent to the slide was not bound and was lost.

**Figure 4.**
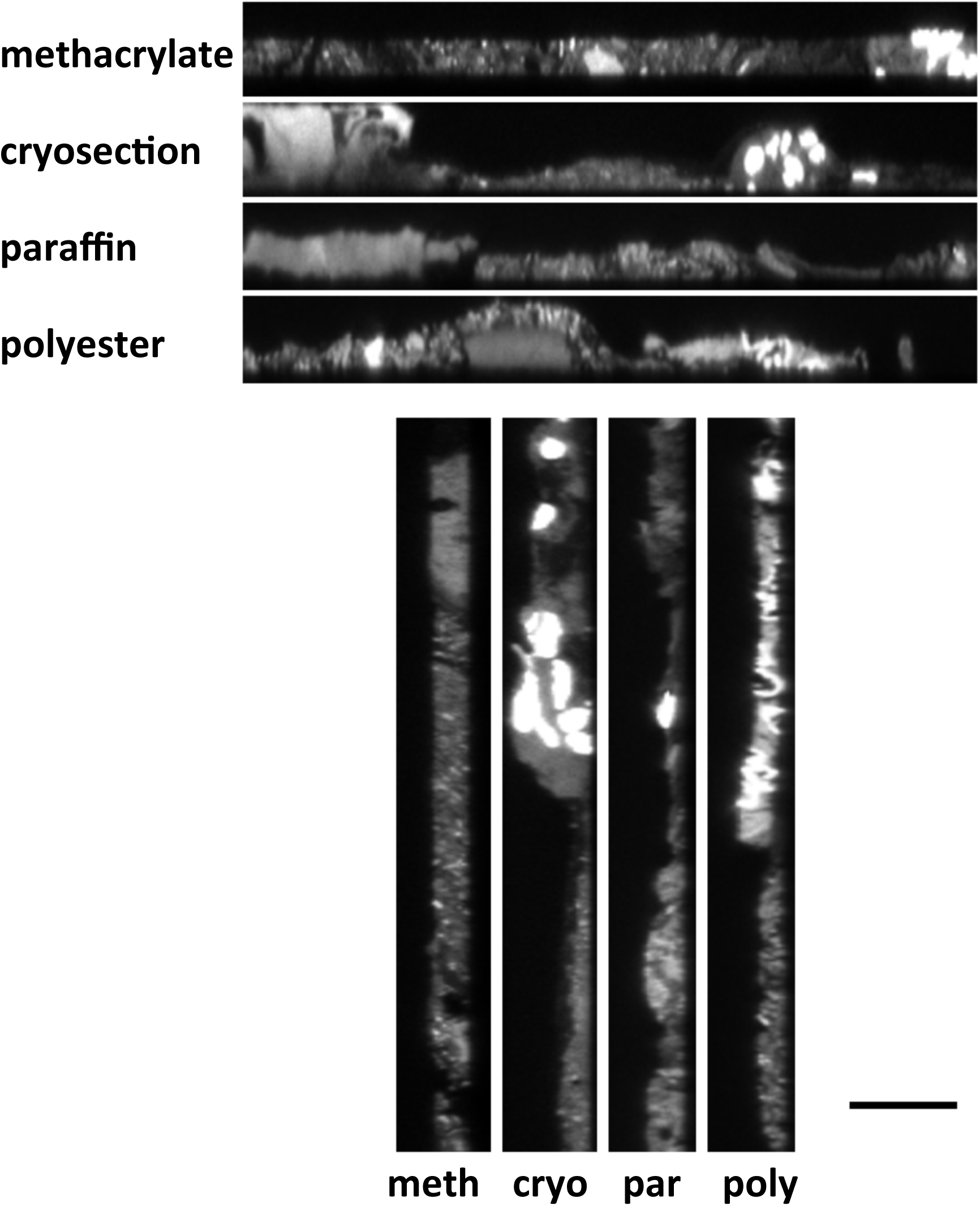
Three-dimensional structure of the gut microbial community is preserved by embedding in methacrylate resin. Projections along the *xz*- and *yz*-planes show that the methacrylate-embedded section is of consistent thickness throughout, whereas the microbial community collapses onto the slide or is partially washed away when using cryosectioning, paraffin, and polyester embedding. From a stack of 35 images at half-micron intervals forming a *z*-series for each section, the projections of the stack along the *xz*-plane (top) and *yz*-plane (boYom) are shown. The positions of the *xz-* and *yz*-images in the *xy*-plane are shown by the magenta lines in Figure 3. Scale bar = 20 microns.

**Figure 3.**
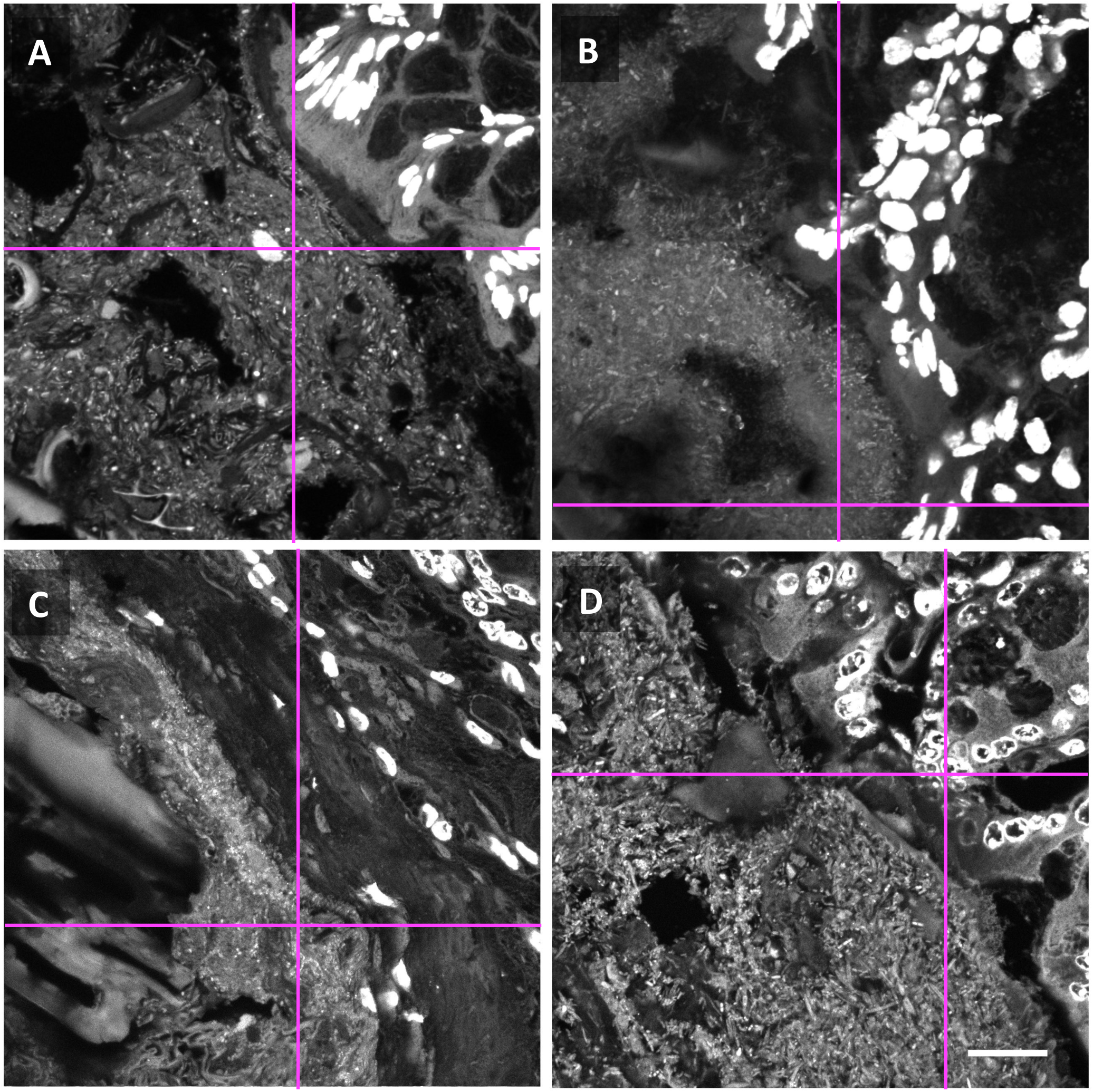
Multiple preparation methods permit imaging of the gut microbial community. Mouse proximal colon embedded in methacrylate resin (A), cryosectioned (B), or processed by embedding and sectioning in paraffin (C) or polyester wax (D). Sections were stained with DAPI and washed, and images were acquired with a confocal microscope using 405 nm excitation. Mucosal tissue (upper right in each panel) shows bright DAPI staining in cell nuclei. Luminal contents including microbes and ingested food particles are at lower left in each image. Magenta lines indicate positions of projections shown in Figure 4. Scale bar = 20 microns.

The differences in results from different embedment materials can be understood in terms of their hydrophobic versus hydrophilic character. Both paraffin and polyester embedments, being hydrophobic, require organic solvents to remove the embedding wax after sectioning before staining or probe hybridization. Redistribution, collapse, or loss of luminal contents could have occurred after removal of the embedding wax, during passage from the organic solvent to the aqueous solution needed for hybridization or during subsequent washing steps. In contrast, Technovit H8100 is a covalently cross-linked methacrylate resin. It is hydrophilic and allows probe hybridization after sectioning without removal of the embedding resin. In the case of cryosectioning, the OCT compound only works to immobilize contents while at low temperature. After cryosectioning, the necessity to warm the section before exposing it to an aqueous solution or to a fixative essentially negated the OCT immobilization. These treatments removed the OCT compound by dilution and had to be carried out at elevated temperature (0-4°C), conditions under which redistribution and collapse could occur. In summary, of the embedments examined, only Technovit H8100 enabled both retention and visualization of the three-dimensional distribution of bacteria and food particles in the intestinal lumen.

Technovit H8100 also proved to be a very stable embedment. Blocks or sections could be stored for long periods (at least 2 years) and probed at a later time, producing results indistinguishable from those obtained by probing immediately after sectioning. Also, probed sections were stable and could be imaged at a later date without noticeable differences except those caused by photobleaching during the initial image acquisition. Consequently, all further experiments were carried out on Technovit H8100 embedded and sectioned material.

### Comparison of fixation methods for preservation and visualization of mucus

Host-derived mucins play an important role in determining the composition and biogeography of microbes in the gut. Hence, it is important to be able to visualize mucus as well as to probe for microbes. Historically, Carnoy’s fixative has been used for the fixation of intestinal samples to preserve mucus produced by the host tissues (Johansson et al. 2008, Hansson & Johansson 2010). However, the fixative commonly used in the established microbial FISH protocol (Moter & Göbel 2000; Amann & Fuchs 2008) is PFA. Consequently, we evaluated preservation of overall mucus structure by fixation with PFA compared with Carnoy’s fixative. Freshly dissected or previously frozen intestinal samples were fixed in parallel either with Carnoy’s fixative or in PFA, embedded in Technovit H8100, sectioned, probed in parallel and imaged. Mucus was visualized with a fluorescent WGA which binds to sialic acid residues known to be abundant in colonic mucin (Rhodes 1989, Matsuo et al. 1997) and by indirect immunofluorescence using an anti-MCM as the primary antibody. The results (Figs 5 and 6) showed that both fixation procedures were effective in preserving and allowing the visualization of intestinal mucus. Mucus in the lumen, lining the epithelial boundary and in goblet cells all could be visualized. No significant differences were detected between Carnoy’s fixative and PFA fixation. Mucus staining patterns observed in these experiments were similar to those described in studies that used Carnoy’s fixative to preserve mucus in intestinal sections (Johansson et al. 2008, Hansson & Johansson 2010, Johansson et al. 2011). These results indicate that mucus components stainable with WGA and anti-MCM antibody were preserved in intestinal sections fixed with PFA.

**Figure 6.**
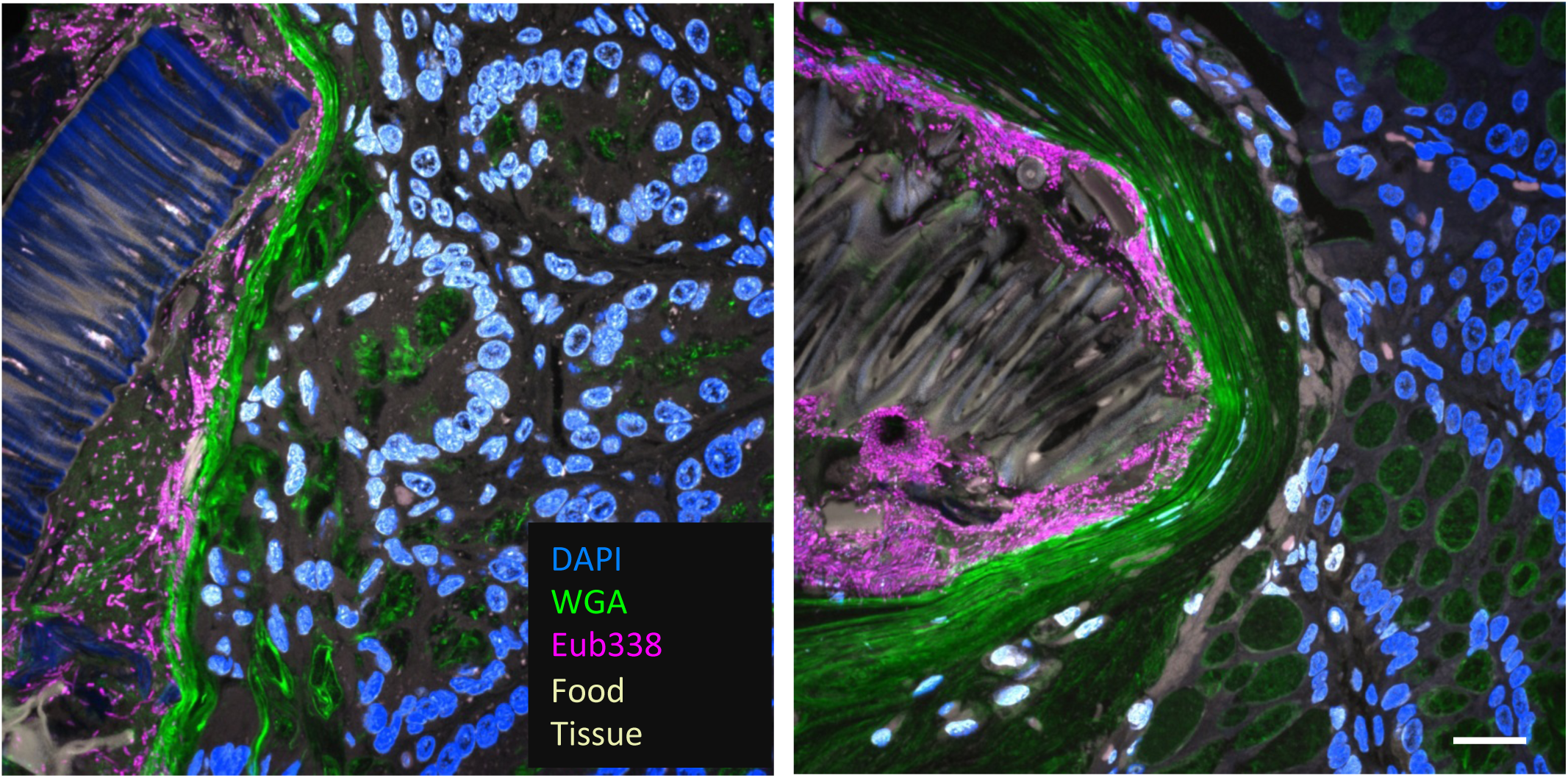
Simultaneous visualization of mucus, FISH-reactive microbes, host tissue, and ingested food with Carnoy fixation and with PFA fixation. Adjacent 1 cm segments of intestine were fixed in Carnoy or PFA, embedded in methacrylate, sectioned, probed in parallel and imaged with identical imaging settings. LeS: Carnoy; right: PFA. Maximum intensity projection of 4 to 5 planes. DAPI stains nucleic acids, wheat germ agglutinin (WGA) stains mucus, and the Eub338 probe hybridizes with most bacteria. Scale bar = 20 microns.

**Figure 5.**
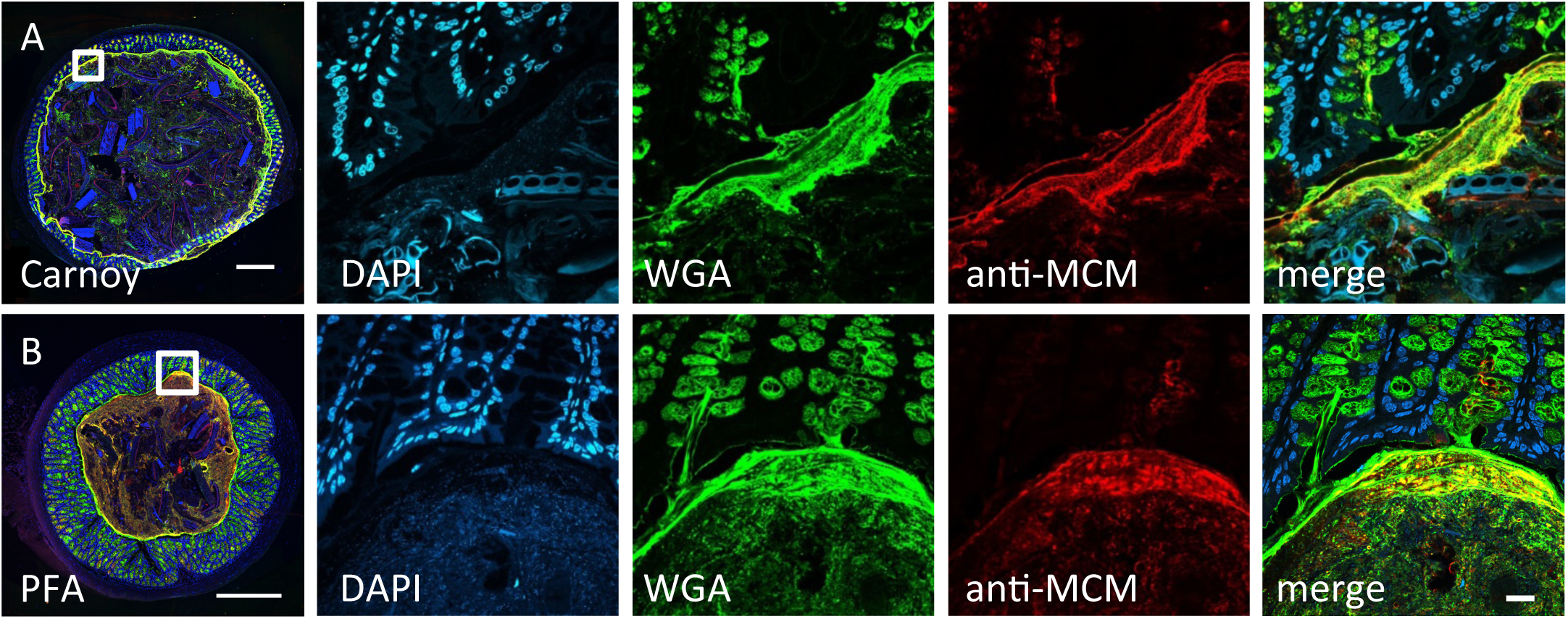
Carnoy and PFA fixation are both capable of preserving mucus, host tissue, and ingested food in intestinal segments that are embedded in methacrylate and sectioned. Overview images (at left) show entire cross-sections of mouse proximal colon fixed with Carnoy (A) or paraformaldehyde (B). Middle panels show higher-magnification images stained with DAPI (blue), wheat germ agglutinin (green) and an antibody against mouse colonic mucin (anti-MCM, red) with a merged overlay image shown at right. Mucus, stained by both wheat germ agglutinin and anti-MCM, is preserved with both fixation methods. Scale bar = 500 microns in overview images, 20 microns in other panels.

### FISH probe reactivity in methacrylate sections with both Carnoy and PFA fixation

FISH is a labeling technique adapted to visualize microbial cells and is the method of choice for studies of microbial spatial structure (Amann & Fuchs, 2008). Importantly, the method provides the opportunity to link taxonomic information of labeled cells to their localization patterns as the oligonucleotide probes used in FISH are designed against taxon-specific regions of the rRNA. We applied a standard FISH protocol to label microbial cells in Technovit H8100-embedded intestinal sections. Specifically, a fluorescently labeled oligonucleotide probe (Eub338-Rhodamine Red X) designed to target most bacteria was used to demonstrate the applicability of the FISH protocol after either Carnoy or PFA fixation. Our results (Fig 6) showed that fluorescent signals from FISH probe-labeled bacterial cells after PFA fixation were comparable to those after Carnoy fixation and that the FISH procedure was also compatible with visualization of mucus.

### Application of the improved protocol to gnotobiotic mouse gut

To illustrate the utility of the optimized embedment protocol, we applied it to gnotobiotic mouse gut colonized by two human gut taxa, one from the phylum Bacteroidetes and one from the phylum Firmicutes. Sample *xy*-plane images are shown in Fig 7. Inspection of the images shows clear differentiation of both of the bacterial taxa as well as food particles, mucus, and host cells. WGA-stained mucus was clearly visible both in goblet cells (Fig 7A) and at the border between mucosa and lumen (Fig 7B). Both of the microbial taxa were found in close proximity to one another as well as to mucus (Fig 7B) and ingested food particles (Fig 7C,D). The food particles themselves ranged widely in size, shape, and autofluorescence spectrum (Fig 7B,C,D) and included both uncolonized cavities and cavities colonized by a dense microbial community (Fig 7D). The variety and complexity of these features presents both challenges and opportunities for the analysis of gut microbial spatial distribution. A quantitative analysis of the distribution of microbes relative to these landmarks in the gnotobiotic gut is presented elsewhere (Mark Welch et al., submitted).

**Figure 7.**
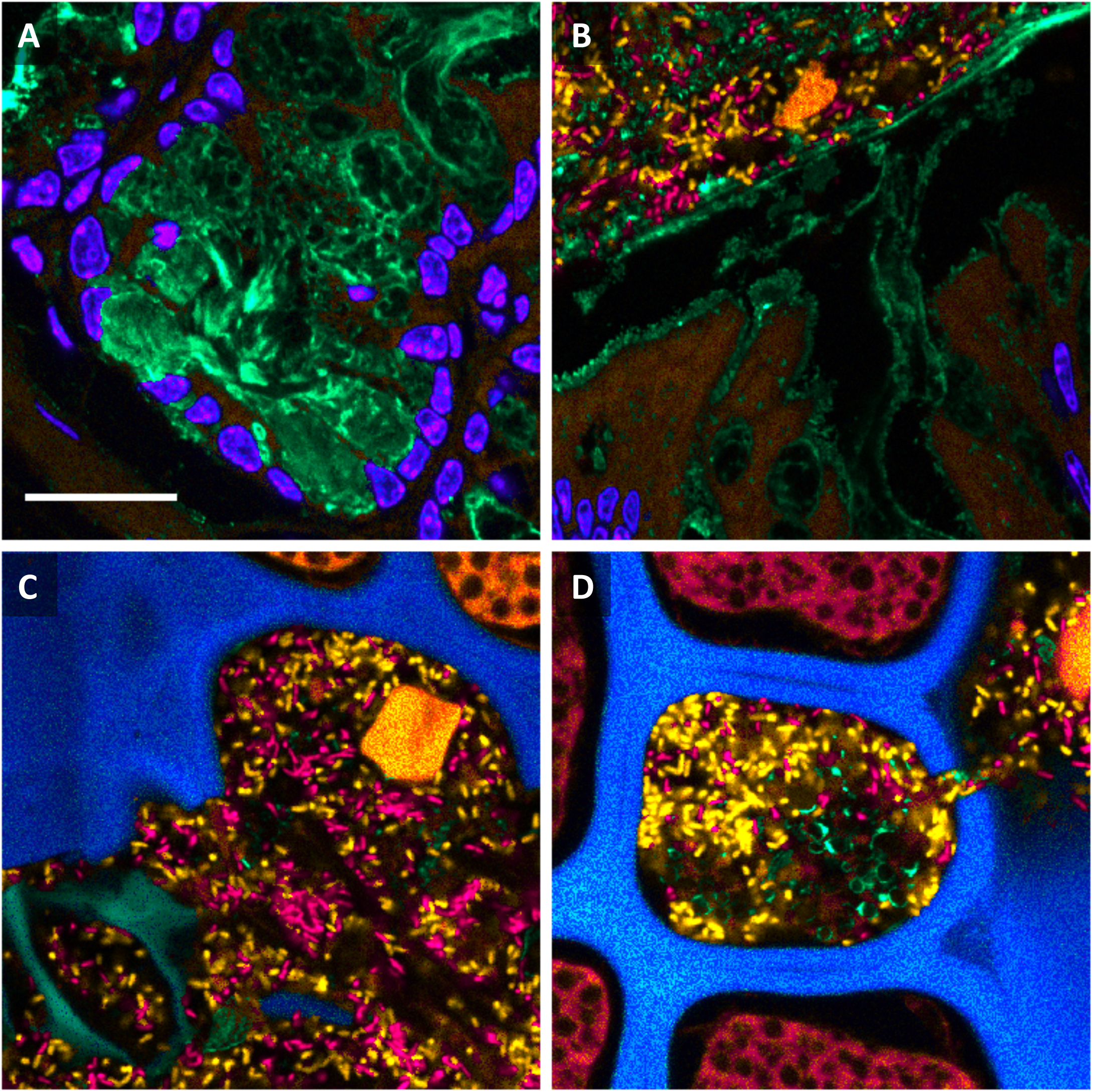
Visualization of the spatial organization of the gut environment using PFA fixation and methacrylate embedment. A gnotobiotic mouse colonized with *B. thetaiotaomicron* and *E. rectale* was fixed with paraformaldehyde, embedded in methacrylate, sectioned, hybridized with oligonucleotide probes to differentiate the two microbial taxa, and stained with DAPI and with fluorophore-labeled wheat germ agglutinin. Fluorescence spectral images were coded to approximate true color. (A): host cell nuclei (blue) and mucus in goblet cells (green). (B): the border between mucosa and lumen. Host cell nuclei are blue, mucus is green*, B. theta* is orange and *E. rectale* is red. Host tissue and a ten-micron-long food particle fluoresce orange. (C): a region of the lumen showing food particles with varying shapes and autofluorescence spectra in blue, green, and orange, signifying different types of food particles. (D): one such food particle possessed both colonized and un-colonized cavities. Scale bar = 20 microns.

## Discussion

A major finding of this study is that special precautions must be taken to preserve the three-dimensional organization of the gut luminal contents including microbes, food particles and mucus. This is not surprising because the luminal contents are not attached to the gut wall and are therefore subject to loss or redistribution. Our results indicate that the protocols involving commonly used embedment media, Tissue-Tek OCT compound, paraffin, or polyester waxes allowed the luminal contents to become redistributed. The reasons for the redistribution seem evident. In the case of the OCT compound, warming of the cryosections permits a loss of immobilization prior to chemical fixation. In the case of paraffin or polyester embedments, their hydrophobicity requires extraction with organic solvents prior to FISH labeling which permits redistribution to occur during and after the extraction step. A cautionary note is that the redistribution was only evident by confocal examination through the depth of the sections. Casual examination of two-dimensional images was not sufficient to reveal the redistribution.

We found that only the hydrophilic, cross-linked methacrylate resin, Technovit H8100 allowed preservation of three-dimensional organization. It was compatible with pre-embedment fixation and post-sectioning labeling. With this embedment, we demonstrated that the micron-scale, three-dimensional spatial arrangement of microbial cells, food particles and mucus could be preserved and visualized in intestinal sections. In addition, we demonstrated that the Technovit H8100 embedment was compatible with two different types of mucus labeling methods, immunostaining with anti-MCM antibody and staining with WGA and that these mucus stains could be applied to intestinal sections that were already labeled with FISH probes. Taken together, these methods allow simultaneous visualization of mucus and microbial taxa of interest along with other biogeographical landmarks that emit autofluorescence, including host tissues and food particles in the same field of view.

We showed that when intact gut segments are fixed and embedded in methacrylate, both Carnoy fixation and PFA fixation are capable of preserving morphological features of the sample including microbes, ingested food particles, and mucus. We suggest that investigators choose a fixation method based on pilot experiments to determine which fixative best preserves epitopes, staining properties, or other features of interest in the individual study.

Finally, we applied the optimized protocol to the gnotobiotic mouse gut colonized by two representative human taxa and illustrated how spatial information at the micron scale could be observed. Appreciation is growing for the importance of spatial organization to understanding microbiome function. The protocol optimized in this study should be useful for studies of microbial spatial organization both among members of the microbial community as well as in relation to a variety of host biogeographical landmarks. The protocol will likely be of value not only for studies of tissue such as gut with non-adherent luminal contents, but for other samples as well in which microbe-microbe and host-microbe relationships are important.

## Acknowledgements

We thank Louie Kerr for microscopy support, Alan Kuzirian and George Bell for advice, and Braden Tierney and Jake Casper for sectioning and imaging technical assistance. We are deeply grateful to Nathan McNulty and Jeffrey I. Gordon for generously providing samples from the two-taxon gnotobiotic mice, and to Shanta Messerli for generously providing samples from conventional mice. This work was supported by the U.S. National Science Foundation grant 1650141 and by NIH National Institute of Dental and Craniofacial Research grant DE022586.

